# Tamoxifen, associated to the conservative CKD treatment, promoted additional antifibrotic effect on experimental hypertensive nephrosclerosis

**DOI:** 10.1101/2022.10.13.512065

**Authors:** Camilla Fanelli, Felipe M Ornellas, Giovanna A Celestrino, Danielly N Carmagnani, Ana LR Francini, Irene L Noronha

**Author notes:** Corresponding Author:* Camilla Fanelli, PhD, Laboratório de Nefrologia Celular, Genética e Molecular, Faculdade de Medicina da Universidade de São Paulo, Av. Dr. Arnaldo, 455, 4° andar, Lab 4304, CEP 01246-903, São Paulo, Brasil.

## Abstract

CKD progression depends on the activation of an intricate set of hemodynamic and inflammatory mechanisms, promoting renal leukocyte infiltration, inflammation and fibrosis, leading to renal function loss. There are currently no specific drugs to detain fibrogenesis, which is a common end-point for different nephropathies. Clinical therapy for CKD is mostly based on the management of hypertension and proteinuria, partially achieved with renin-angiotensin-aldosterone system (RAAS) blockers, and the control of inflammation by immunosuppressive drugs. The aim of the present study was to verify if the administration of tamoxifen (TAM), an estrogen receptor modulator, clinically employed in the treatment of breast cancer and predicted to exert antifibrotic effects, would promote additional benefits when associated to a currently used therapeutic scheme for the conservative management of experimental CKD. Wistar rats underwent the NAME model of hypertensive nephrosclerosis, obtained by daily oral administration of a nitric oxide synthesis inhibitor, associated to dietary sodium overload. The therapeutic association of TAM to losartan (LOS), and mofetil mycophenolate (MMF) effectively reduced the severe hypertension, marked albuminuria and glomerular damage exhibited by NAME animals. Moreover, the association also succeeded in limiting renal inflammation in this model, and promoted further reduction of ECM interstitial accumulation and renal fibrosis, compared to the monotherapies. According to our results, the association of TAM to the currently used conservative treatment of CKD added significant antifibrotic effects both *in vivo* and *in vitro*, and may represent an alternative to slow the progression of chronic nephropathy.

## INTRODUCTION

The pathogenesis of CKD involves an intricate process of both hemodynamic and inflammatory mechanisms that leads to renal fibrosis and progressive loss of function. Glomerular and systemic hypertension, increased production of cytokines and growth factors, renal infiltration of inflammatory cells, inordinate fibroblast proliferation and transdifferentiation into myofibroblasts have been described in different human nephropathies and experimental CKD models [1–4].

It is widely known that overactivation of both systemic and intrarenal renin-angiotensin-aldosterone system (RAAS) contributes to the progression of CKD [5,6]. Once bound to its specific receptor (AT1), the active peptide Angiotensin II (AII) promotes renal and systemic vasoconstriction and tubular sodium conservation, leading to the elevation of blood pressure [1,2,6]. Moreover, AII also exerts proinflammatory effects, once it stimulates cell proliferation, fibroblast activation and further accumulation of extracellular matrix (ECM). Actually, since the discovery of the angiotensin ii converting enzyme inhibitors (ACEi) and the AT1 receptor blockers (ARB), such as losartan (LOS), RAAS suppression remains to be the best option available to slow the progression of CKD, although this strategy does not fully halt the progression of renal fibrosis and loss of function [7–10].

It is well known that inflammation and increased ECM production exert an important pathogenic role to the development and progression of CKD, regardless of its etiology. Accordingly, we have previously demonstrated that treatment with anti-inflammatory drugs such as mycophenolate mofetil (MMF), which promotes antiproliferative effects on T-cells, presented effective renoprotection in experimental CKD, in the hypertensive nephrosclerosis model obtained by chronic inhibition of nitric oxide synthesis (NAME model), in the 5/6 nephrectomy (Nx) and in the streptozotocin-induced diabetes [11–13]. Moreover, Nx rats treated with an association of MMF + LOS presented significantly less severe CKD when compared to animals receiving the respective monotherapies [14].

Considering that renal fibrosis with tissue scarring is the final common pathway of CKD, therapeutic interventions with antifibrotic drugs could represent an attractive choice of therapy to arrest fibrogenesis in progressive nephropathies. In this context, tamoxifen (TAM), an estrogen receptor modulator clinically used in the treatment of breast cancer, has been demonstrated to also be effective in treating abnormal healing disorders, such as retroperitoneal fibrosis, sclerosing encapsulated peritonitis, fibrosing mediastinitis, among other fibroproliferative conditions [15–18].

Moreover, we have recently shown that TAM treatment prevented the development of glomerulosclerosis and interstitial expansion in rats submitted to NO inhibition, even having no effects on the marked hypertension characteristic of this model [19].

Motivated by the positive responses achieved by each employed monotherapy in the treatment of progressive nephropathy associated to the NAME model of CKD, in the present study, we sought to verify if the administration of an association of LOS+MMF+TAM could promote additional renoprotection compared to the respective monotherapies, once the mechanisms of action of each drug are different and somehow complementary.

## MATERIAL AND METHODS

### Experimental Groups and Protocol

The experimental model of hypertensive nephrosclerosis employed in the present study was described in our previously published article, entitled “Gender Differences in the Progression of Experimental Chronic Kidney Disease Induced by Chronic Nitric Oxide Inhibition” [4]. Thirty-five male Wistar rats aged between 7 and 8 weeks were kept under controlled temperature (23±1°C), on a 12/12 hours’ light/dark cycle with *ad libitum* access to tap water and HS diet (3.12% Na, Nuvital, Brazil). After 2 weeks of adaptation to HS_diet, 30 of these animals were submitted to the NAME experimental model of chronic nitric oxide (NO) inhibition. This CKD model was obtained by oral daily administration of 70 mg/kg/d of Nω-nitro-L-arginin metil-ester (L-NAME - Sigma Chemical CO, St. Louis, USA), a L-arginine analogue, diluted on drinking water, associated to HS diet. NAME rats were divided among the following 5 groups: **NAME:** Animals submitted to the NAME model and keep untreated; **LOS:** NAME animals treated with 50 mg/Kg/d of losartan (LOS) diluted in drinking water; **MMF:** NAME rats treated with 10 mg/Kg/d of Micofenolate Mofetil (MMF) administered daily by gavage; **TAM:** NAME animals receiving 10 mg/Kg/d of Tamoxifen (TAM) and **LOS+MMF+TAM:** NAME rats treated with LOS, MMF and TAM simultaneously. Five additional animals received only HS and were used as **Control**.

All groups were followed for 30 days. Body weight was monitored weekly and at the end of this period, blood pressure was evaluated by the tail-cuff pressure method, using a noninvasive system (RTBP 2045; Kent Scientific). Additionally, 24-hour urinary albumin excretion rate (24h-UAE) was analyzed by radial immunodiffusion, as described elsewhere [4, 20]. Animals were anesthetized with an intraperitoneal (IP) injection of 60 mg/kg of sodium pentobarbital and submitted to total nephrectomy followed by euthanasia through overdose of sodium pentobarbital, 80 mg/kg IP.

### Histological Analysis

As described elsewhere [4], kidneys obtained from total nephrectomy were cut in two midcoronal renal slices and pre-fixed with Duboscq-Brazil for 30 minutes, followed by 24-hour post-fixation in buffered 4% formaldehyde. Tissue samples were embedded in paraffin, through conventional techniques, renal tissue sections of 4-μm thickness were obtained and submitted to histological analysis for the assessment of glomerular and tubulointerstitial alterations.

The percentage of glomerulosclerosis and collapsed glomeruli were evaluated in periodic Acid-Schiff (PAS) staining samples, through the analysis of at least 50 randomly sampled glomerular tuft profiles per rat. The criteria used to define sclerotic glomeruli was the presence of segmental hyalinosis lesions, usually with adhesion to Bowman’s capsule. Collapsed glomeruli were defined by their reduced size, wrinkling basement membrane and collapsed capillary loops. Interstitial fibrosis was quantitatively evaluated in Masson-stained sections by a point counting technique [4, 21].

### Immunohistochemical Analysis

Immunohistochemistry (IHC) assays were performed to identify interstitial macrophage and T-cell infiltration in renal sections, as well as to evaluate tubulointerstitial cell proliferation and to quantify the percentage of tubulointerstitial area occupied by α-smooth muscle actin (α-SMA), possibly indicating the presence of myofibroblasts in the renal cortex [2], collagen I and fibronectin. A mouse monoclonal anti-ED1 antibody (Serotec, Oxford, UK) was used to identify macrophages through the APAAP (alkaline phosphatase anti-alkaline phosphatase) technique. Monoclonal mouse anti-CD3 (Dako, Glostrup, Denmark) and anti-α-SMA (Sigma Chemical CO, St. Louis, USA) antibodies were used, to identify T-cells and myofibroblasts, respectively, through a streptavidin-biotin-alkaline phosphatase (Strep-AP) IHC technique. In both APAAP and Strep-AP techniques, the reactions were developed with a fast-red dye solution. Tubulointerstitial cell proliferation was detected by a monoclonal mouse anti-PCNA antibody (Dako, Glostrup, Denmark), while collagen I and fibronectin positivity were detected with polyclonal anti-collagen I (Rockland Immunochemicals, Inc., NY, USA) and anti-fibronectin (Sigma Chemical CO, St. Louis, USA) primary antibodies, using a streptavidin-biotin-horseradish peroxidase (Strep-HRP) IHC technique. Samples were developed with a DAB dye solution.

### In vitro Experiments

In order to verify the specific antifibrotic activity of TAM alone, and also to investigate if this activity would be somehow inhibited or impaired by the association with other drugs, we performed cell culture experiments using a rat renal fibroblast cell line (NRK-49F; American Type Culture Collection, Manassas, VA). For this purpose, 1×10^5^ NRK-49F cells were cultured under 37°C and 5% CO_2_ in plastic culture plates with Dulbecco’s Modified Eagle Medium (DMEM-Low glucose, Invitrogen, USA) containing 5% inactivated fetal bovine serum (FBS; Gibco, Carlsbad, MO, USA), 100 units/mL penicillin, and 100 mg/mL streptomycin antibiotic solution (Gibco). Once cells reached 80% of confluence, the culture medium was replaced by DMEM-Low with 100 units/mL penicillin, and 100 mg/mL streptomycin antibiotic solution, plus the specific stimuli, as follows; <u>Control</u>, NRK-49 cells receiving no additional stimuli or treatment diluted in the culture medium, <u>IL-1β+AngII</u>, NRK-49 cells whose culture medium was supplemented with 400pg/mL of recombinant human IL-1β (PeproTech, Cranbury, NJ, USA) and 1×10^-7^M Angiotensin II acetate human (Sigma), LOS, IL-1β+AngII cells whose culture medium was further supplemented with 10 μM of Losartan, <u>TAM</u>, IL-1β+AngII cells whose culture medium was further supplemented with 5μM of Tamoxifen citrate (Sigma) and <u>LOS+TAM</u>, NRK-49 cells receiving all the above mentioned supplements. Cells were kept under the described treatments for 24h.

### Immunocytochemistry

Immunocytochemistry (ICC) assays were performed to characterize the constitutive expression of vimentin in NRK-49 cells, using a mouse monoclonal anti-vimentin primary antibody (Sigma Chemical CO, St. Louis, USA). The activation of fibroblasts after IL-1β+ AngII stimulus, was evaluated through the positivity of these cells for α-SMA, with a mouse monoclonal anti-α-SMA antibody (Sigma Chemical CO, St. Louis, USA). Moreover, ICC was also employed to analyze the expression of collagen I and fibronectin in NRK-49 cells submitted to the different treatments, employing, respectively, the rabbit polyclonal anti-collagen I (Rockland Immunochemicals, Inc., NY, USA) and anti-fibronectin (Sigma Chemical CO, St. Louis, USA) primary antibodies. Vimentin, collagen I and fibronectin ICC were performed through a streptavidin-biotin-alkaline phosphatase (Strep-AP) technique. Reactions were developed with a fast-red dye solution. α-SMA was detected through a streptavidin-biotin-horseradish peroxidase (Strep-HRP) ICC technique, developed with a DAB dye solution.

### Real time RT-PCR

Quantitative real-time polymerase chain reaction (PCR) of cultured NRK-49F cells was performed to measure the relative gene expression of TGFβ, SMAD3, SMAD7, Collagen type I, collagen type III and Fibronectin, using Actinβ as a housekeeping control, as previously described. Total NRK-49F cells RNA extraction was carried out with RNeasy Plus Kit (Qiagen, MD, EUA), following the instructions of the manufacturer. Reverse transcription (RT) was performed with M-MLV enzyme kit (Promega) and qPCR was conducted with the Syber GreenER qPCR Super Mix Universal (Invitrogen), in the StepOne Plus equipment (Applied Biosystetems - Life Technologies). Quantitative comparisons were obtained using the ΔΔCT method (Applied Biosystems, Singapore, Singapore). Primer sequences for amplifying target genes were: Tgfb1 NM_021578.2, left primer: GCTGAACCAAGGAGACGGAA, right primer: CATGAGGAGCAGGAAGGGTC, Smad3 NM_013095.3 left primer: GAGACATTCCACGCTTCACA, right primer: AAAGACCTCCCCTCCAATGT, Smad7 NM_030858.2 left primer: TCTCCCCCTCCTCCTTACTC, right primer: CAGGCTCCAGAAGAAGTTGG, Coll1a1 NM_053304.1 left primer: AGCTGGTGCTAAGGGTGAAG, right primer: GCAATACCAGGAGCACCATT, Coll3a1 NM_053304.1 left primer: AGCTGGTGCTAAGGGTGAAG, right primer: GCAATACCAGGAGCACCATT, Fn1 NM_019143.2 left primer CTCCCGGAACAGATGCAATG, right primer ATCCAGCTGAAGCACTCTGT and Actb NM_031144.3 left primer: AGGGAAATCGTGCGTGACAT, right primer: CCATACCCAGGAAGGAAGGC.

### Statistical Analysis

Results were presented as mean ± SEM. Differences among all groups were analyzed by one-way ANOVA with Dunnet’s multiple comparison post-test. Means were considered statistically different when p<0.05 [22]. All statistical analyses were realized using the Graph-Pad Prism^™^ 5.01 software.

## RESULTS

### Association of LOS+MMF+TAM was effective in reducing blood pressure and albuminuria

As expected, rats receiving L-NAME exhibited severe hypertension when compared to Control (213 ± 5 vs. 130 ± 3 mmHg). LOS or MMF as monotherapies, as well as the association of LOS+MMF+TAM promoted reduction in the blood pressure levels (183 ± 12, 173 ± 9 and 173 ± 3 mmHg, respectively), as shown in **Figure 1A**. NAME animals also developed marked albuminuria (147.3 ±30.5 vs. 1.1 ± 0.4 mg/24h in the Control group), which was significantly reduced with all the employed monotherapies; LOS (14.2 ± 4.4 mg/24h), MMF (11.1 ± 7.8 mg/24h) and TAM (24.1 ± 4.6 mg/24h). It is worth mentioning that the treatment with the association of LOS+MMF+TAM promoted the most prominent reduction of this parameter, reaching values similar to the control group (4.1 ± 1.3 mg/24h), as presented in **Figure 1B**.

**Figure 1.**
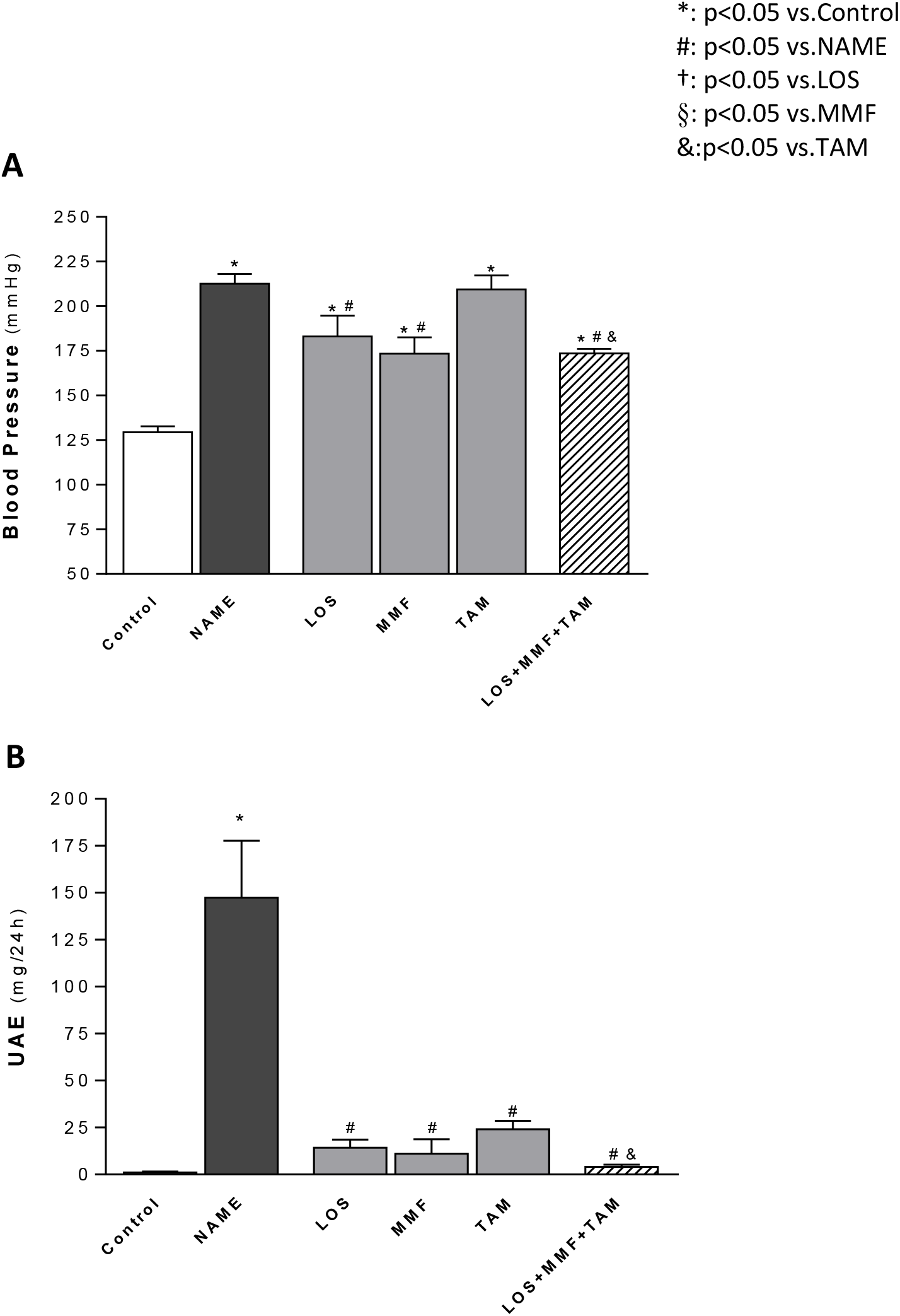
Bar graph representation of blood pressure, measured by tail cuff pressure, (mmHg) **(A)** and urinary albumin excretion rate (UAE, mg/24h) **(B)** in the Control, NAME, LOS, MMF, TAM and LOS+MMF+TAM groups.

### Treatment with the association of LOS+MMF+TAM averted the development of glomerulosclerosis and glomerular collapse in NAME rats

Glomerular structural alterations, characterized by the development of glomerulosclerosis and by the presence of collapsed glomeruli, were accessed by PAS staining. Illustrative micrographs of samples of each experimental group are represented in **Figure 2**. Untreated NAME rats exhibited prominent structural alterations, which were averted by all the employed therapeutic schemes. Bar graphs of the respective quantitative analysis of the percentage of sclerotic and collapsed glomeruli are shown in **Figure 3**. Accordingly, NAME animals shown significant glomerulosclerosis and glomerular collapse, compared to Control rats (2.6 ± 0.7 and 17.3 ± 3.0 vs. 0.3 ± 0.3 and 1.6 ± 0.5 %, respectively). LOS, MMF and TAM monotherapies at least partially averted the development of both glomerulosclerosis and glomerular collapse in NAME rats (0.5 ± 0.2 and 4.1 ± 1.1 %, 1.6 ± 0.7 and 1.2 ± 0.8 %, 1.1 ± 0.3 and 5.1 ± 0.7 %), and the association of LOS+MMF+TAM promoted further protection against these glomerular structural alterations (0.4 ± 0.3 and 0.9 ± 0.5 %).

**Figure 2.**
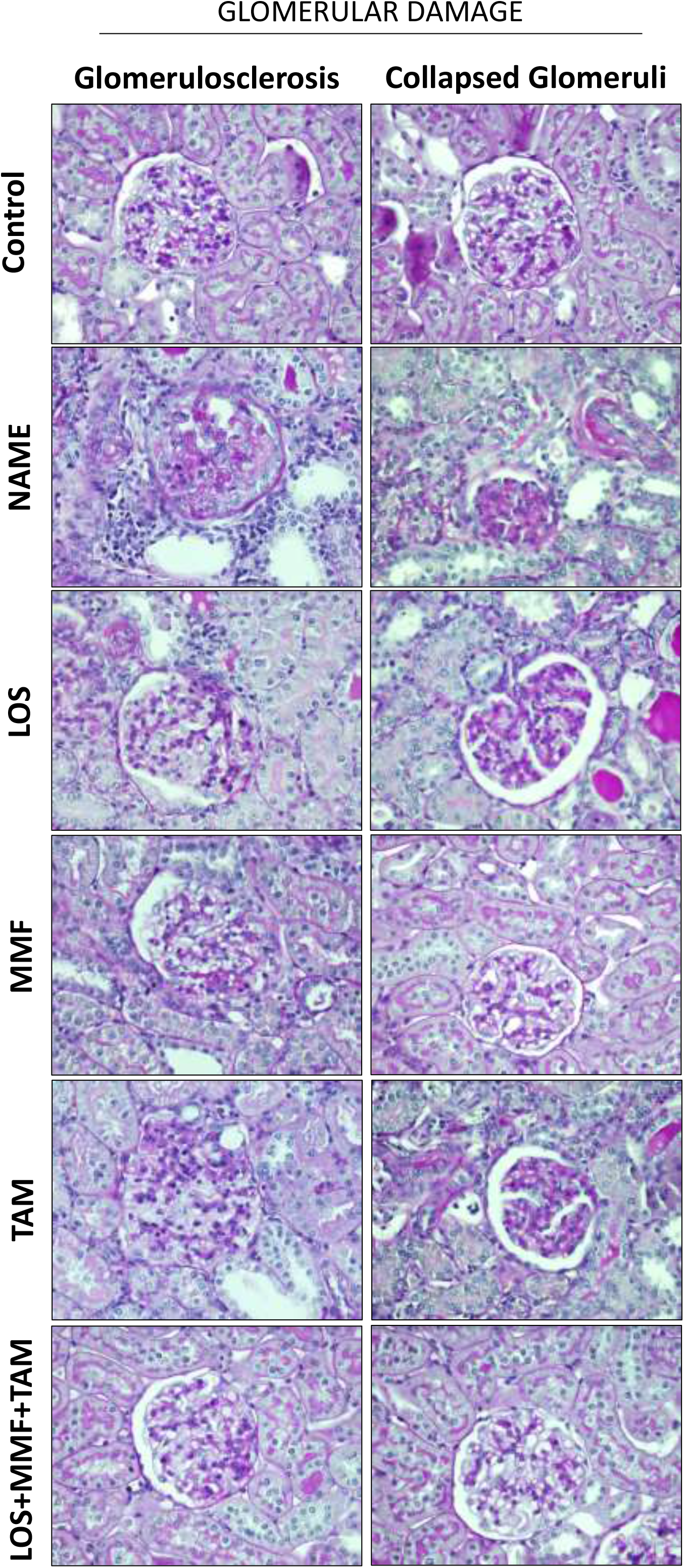
Representative micrographs of glomerulosclerosis **(left)** and collapsed glomeruli **(right)**, accessed in the Control, NAME, LOS, MMF, TAM and LOS+MMF+TAM groups by PAS staining.

**Figure 3.**
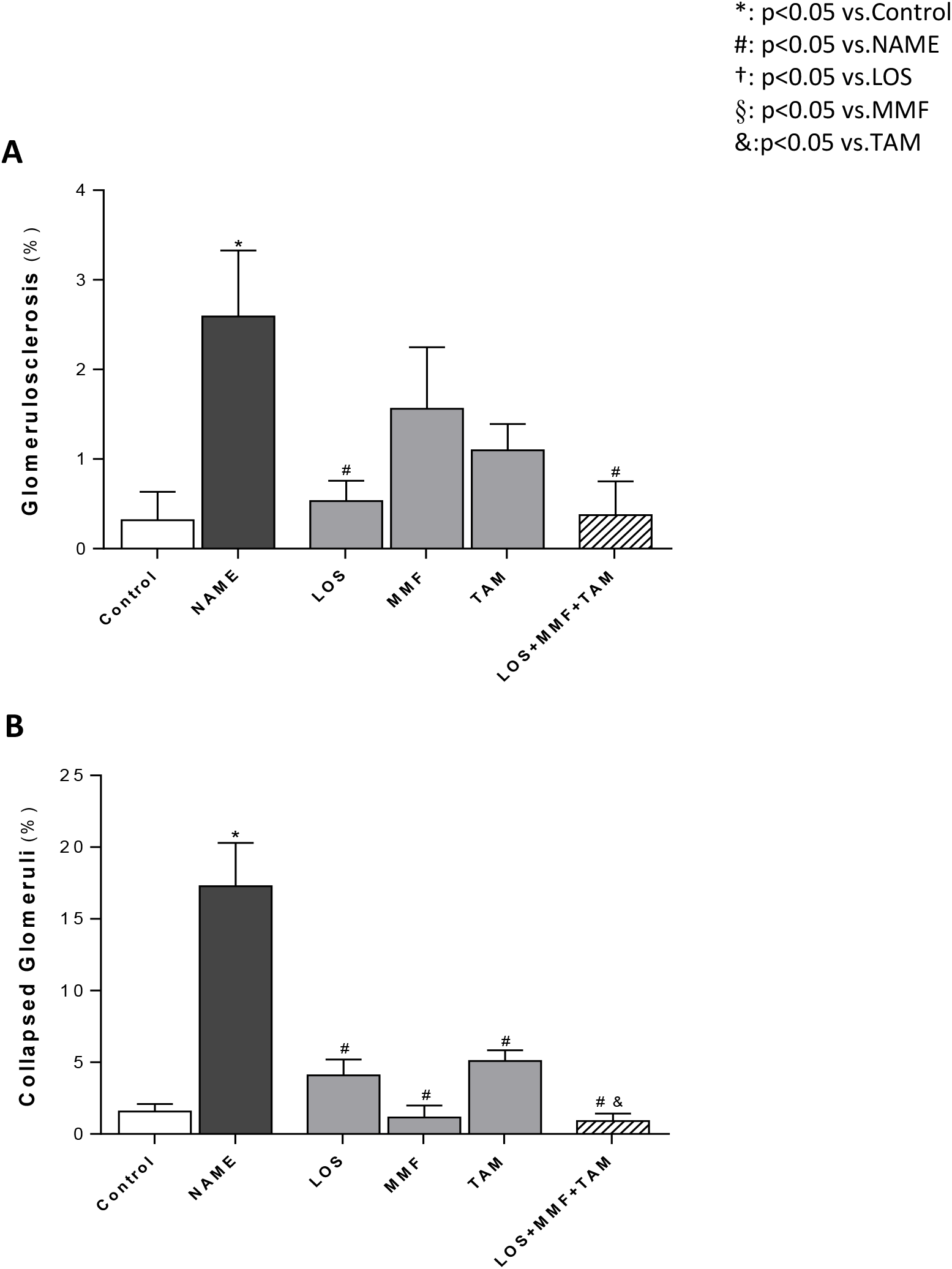
Bar graph representation of the percentage of glomerulosclerosis **(A)** and collapsed glomeruli **(B)** in the Control, NAME, LOS, MMF, TAM and LOS+MMF+TAM groups.

### Combined LOS+MMF+TAM ameliorated interstitial fibrosis in animals submitted to the L-NAME CKD model

As can be seen in **Figure 4**, NAME animals exhibited severe renal interstitial fibrosis, evaluated in Masson trichrome stained renal slides, as well as abundant interstitial α-SMA expression, which indicates the presence of myofibroblasts in the renal cortex. Overexpression of collagen 1 and fibronectin, ECM proteins related to renal fibrosis, was also observed in NAME rats, compared to the Control. Bar graphs of the quantitative analysis of these parameters were presented in **Figure 5**. According to these quantifications, NAME animals exhibited significant renal fibrosis, evidenced by the positivity for Masson staining (1.7 ± 0.3 vs. 0.3 ± 0.1 % in Control), as well as marked α-SMA accumulation (16.2 ± 3.5 vs. 0.4 ± 0.1 % in Control), compared to Control rats. Both Masson positivity and the presence of interstitial myofibroblasts were equally limited by all the employed therapies, both monotherapies with LOS (0.8 ± 0.2 and 5.7 ± 1.0 %), MMF (0.5 ± 0.4 and 7.2 ± 1.0 %) and TAM (0.3 ± 0.1 and 6.4 ± 0.7 %), and the association of drugs (0.4 ± 0.1 and 5.2 ± 1.2 %). Interstitial collagen 1 and fibronectin percentages were also abnormally increased in untreated NAME rats compared to Control animals (21 ± 2 and 16 ±1 vs. 11 ±2 and 7 ± 1 %, respectively). While the interstitial accumulation of collagen 1 was only subtly prevented by TAM monotherapy and the association of LOS+MMF+TAM (16 ± 2 and 17 ± 2 %), interstitial fibronectin percentage was considerably averted by these treatments, especially by the combined therapy (11 ± 1 and 7 ± 2 %).

**Figure 4.**
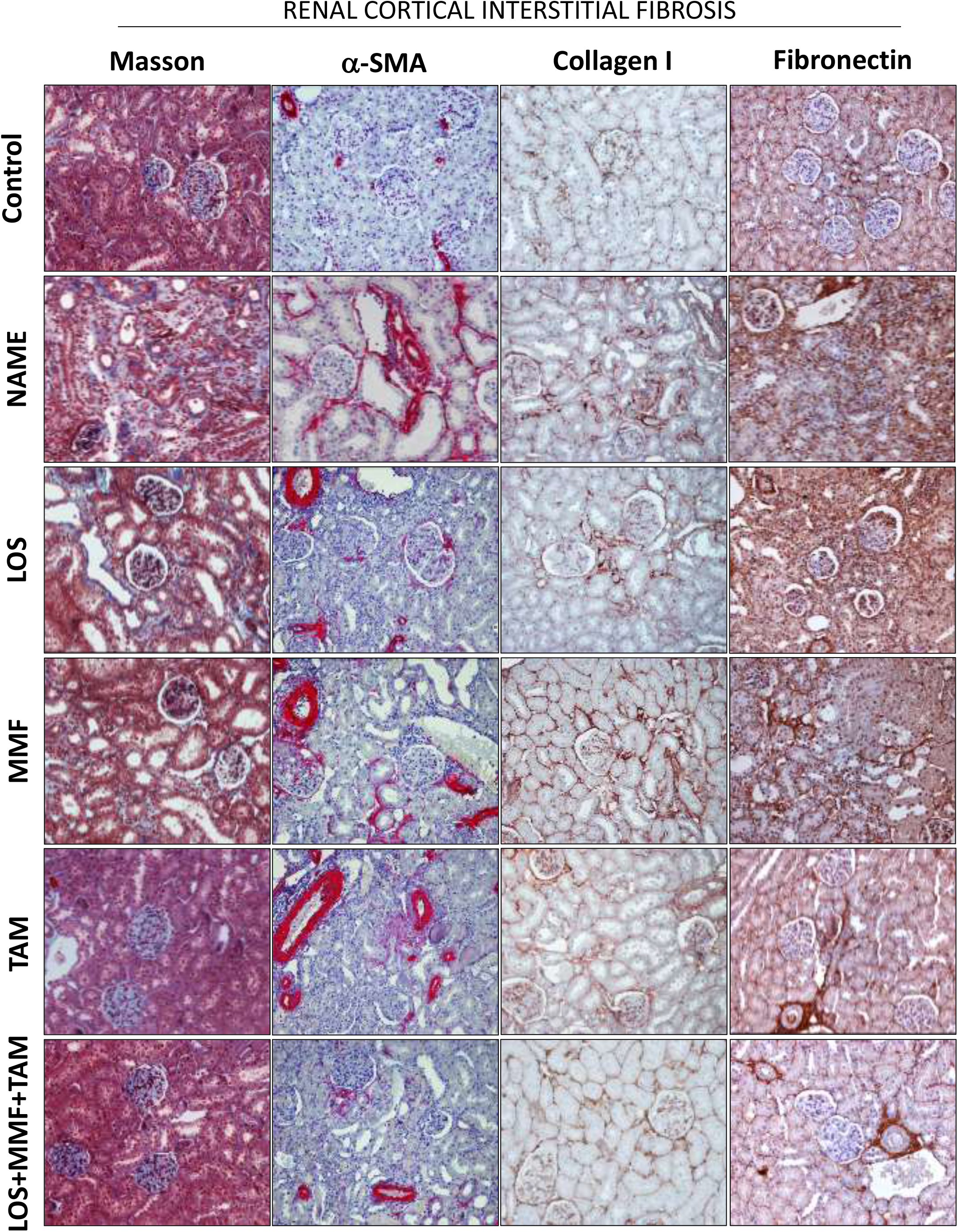
Representative micrographs of renal cortical Interstitial fibrosis, evaluated in Masson’s Trichrome stained slides, interstitial α-SMA, collagen I and fibronectin accumulation, accessed by immunohistochemistry in the Control, NAME, LOS, MMF, TAM and LOS+MMF+TAM groups.

**Figure 5.**
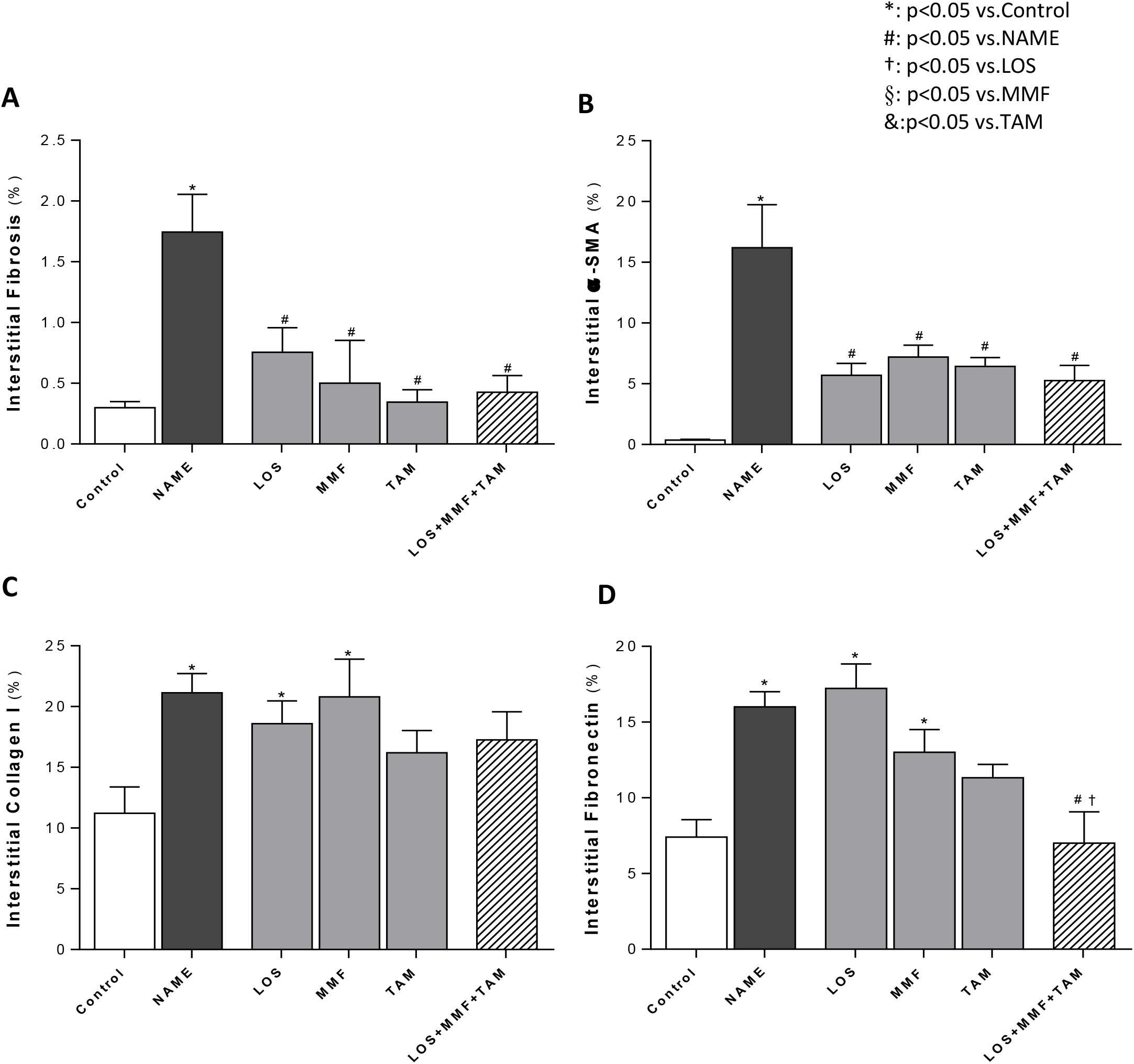
Bar graphs show quantification of percentage of cortical interstitial fibrosis **(A)**, interstitial α-SMA accumulation **(B)**, interstitial collagen I **(C)**, and fibronectin **(D)**, in the Control, NAME, LOS, MMF, TAM and LOS+MMF+TAM groups.

### The association of LOS+MMF+TAM abrogated renal inflammation and reversed interstitial cell proliferation in NAME animals

Illustrative microphotographs of immunohistochemistry for the detection of renal infiltration by macrophages and T-cells and for the evaluation of renal interstitial cell proliferation, in renal sections of animals from each experimental group, can be seen in **Figure 6**. Immunohistochemistry analysis demonstrated that untreated rats submitted to the CKD model induced by L-NAME administration presented marked renal inflammation, characterized by intense cortical infiltration by macrophages and T-lymphocytes, and increased interstitial cell proliferation, evidenced by the presence of interstitial PCNA^+^ cells. The inflammatory cells were detected mainly in the renal interstitium, but were also observed infiltrating glomeruli and in the perivascular area. According to the quantification of these cells, presented in **Figure 7**, untreated NAME rats exhibited significant interstitial macrophage (163 ± 24 cell/mm^2^) and T-cell (127 ± 20 cell/mm^2^) infiltration, compared to the Control (14 ± 5 and 25 ± 5 cell/mm^2^). LOS, MMF and TAM monotherapies statistically reduced these cells (84 ± 17 and 84 ± 11 cell/mm^2^, 50 ± 9 and 42 ± 9 cell/mm^2^ and 82 ± 7 and 38 ± 7 cell/mm^2^, respectively). LOS+MMF+TAM association achieved the numerically lowest values of both macrophage and lymphocyte interstitial infiltration cells (40 ± 6 and 27 ± 5 cell/mm^2^). Similar results were obtained regarding renal cortical cells proliferation. NAME group presented significant increase in PCNA^+^ interstitial cells, compared to Control (142 ± 24 vs. 15 ± 5 cell/mm^2^). All the employed monotherapies reduced cell proliferation in this experimental nephrosclerosis model (LOS: 43 ± 16, MMF: 29 ± 12 and TAM: 31 ± 8 cell/mm^2^), and once more, the lowest number of positive cells were observed with the association of LOS+MMF+TAM (16 ± 6 cell/mm^2^), in which group the interstitial cell proliferation rate was comparable to the observed in the Control.

**Figure 6.**
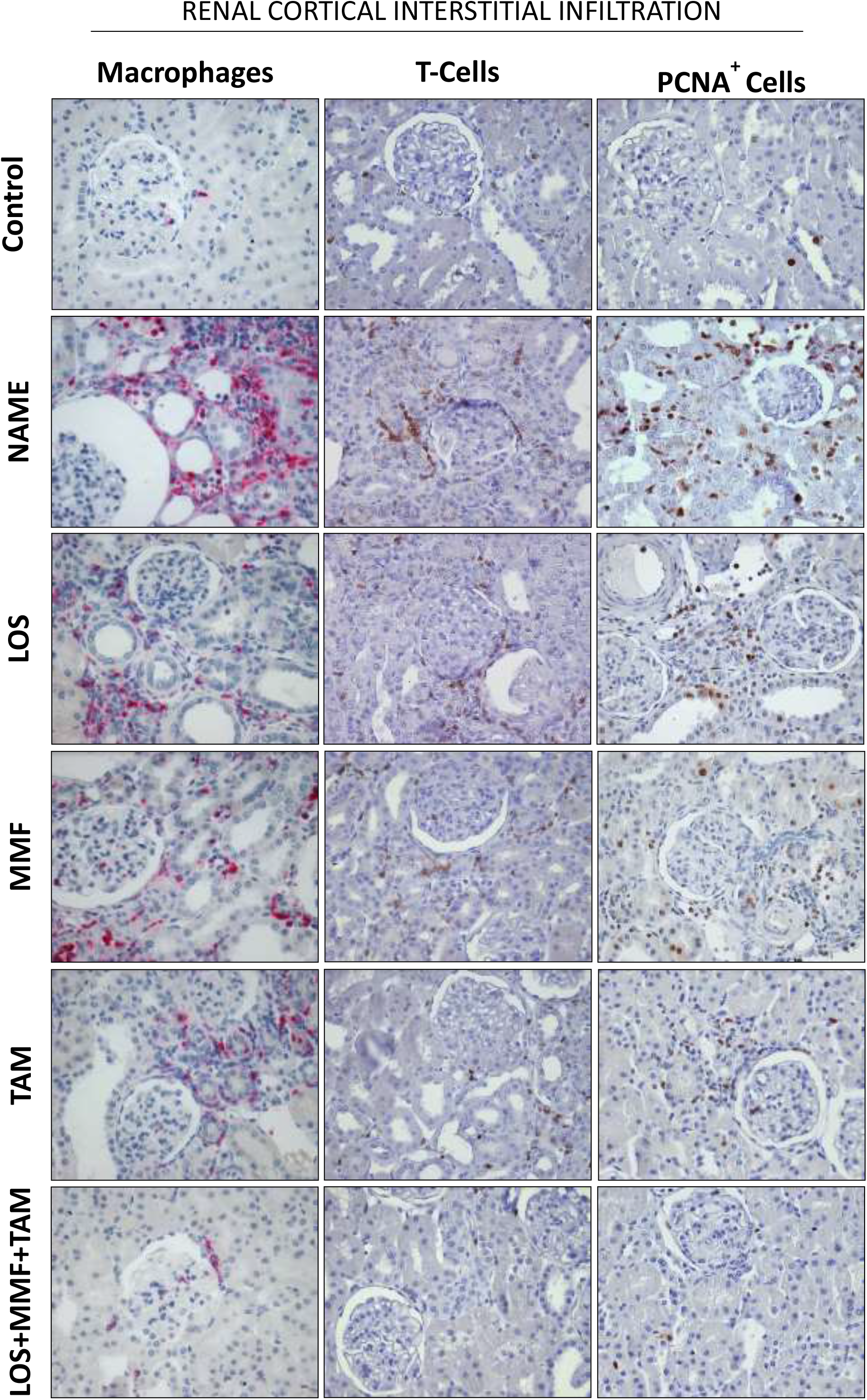
Representative micrographs of renal cortical interstitial infiltration of macrophages, lymphocytes (T-Cells) and PCNA^+^ cells, accessed by immunohistochemistry in the Control, NAME, LOS, MMF, TAM and LOS+MMF+TAM groups.

**Figure 7.**
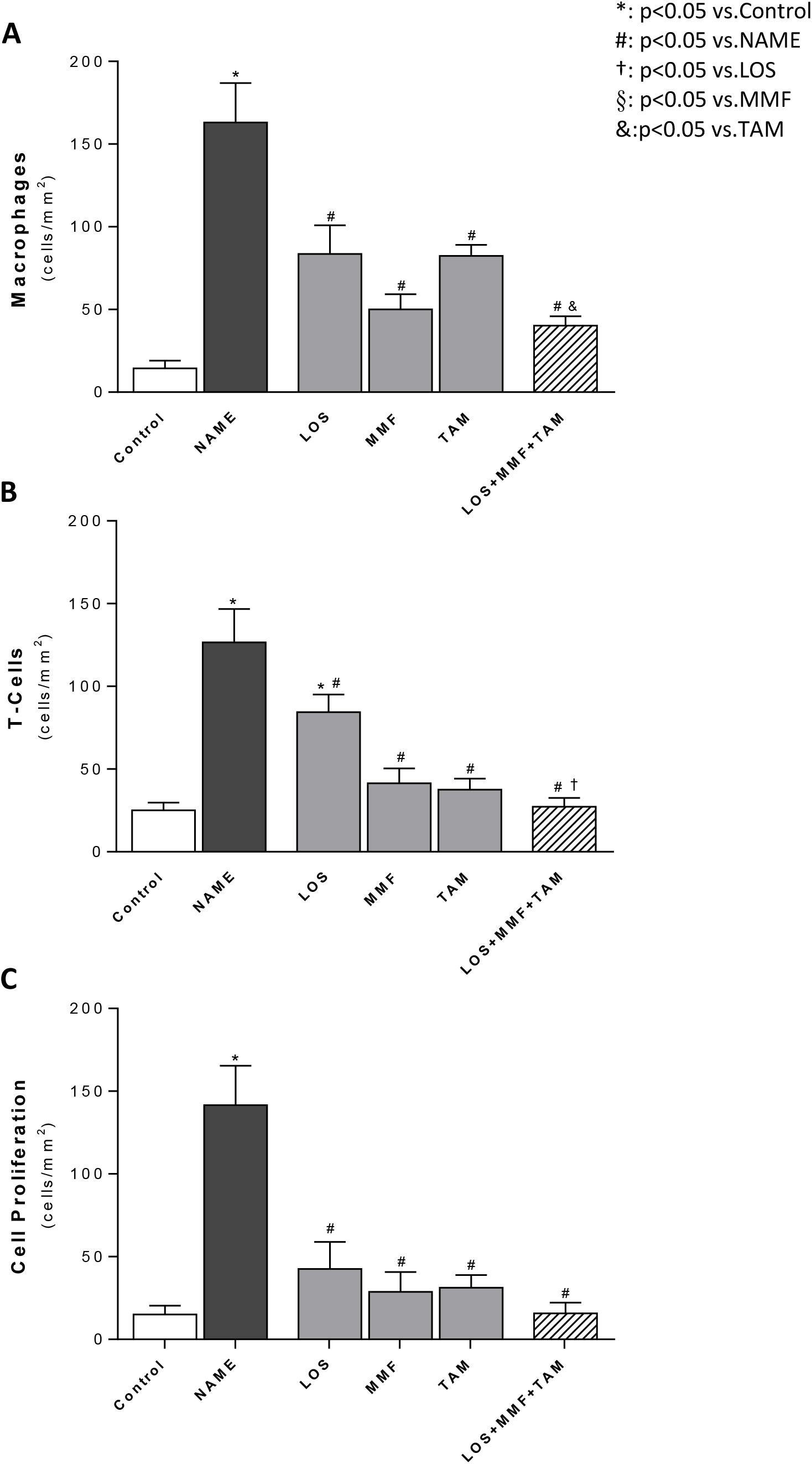
Bar graphs show interstitial infiltration of macrophage **(A)**, T-Cell **(B)** and PCNA^+^ cells **(C)** in the Control, NAME, LOS, MMF, TAM and LOS+MMF+TAM groups.

### TAM in monotherapy or associated to LOS effectively inhibited fibroblasts activation and ECM overproduction in cultured NRK-49F cells

In order to establish an *in vitro* model of activated fibroblasts, which may mimic the subpopulation of renal fibroblasts of our *in vivo* experimental nephrosclerosis model, we stimulate rat renal immortalized fibroblasts from a commercially available cell line (NRK-49F) with IL-1β + AngII. As shown in **Figure 8**, after this stimulus, NRK-49F continued to express vimentin, a cytoskeleton type III intermediate filament, constitutively present in both fibroblasts and myofibroblasts, but also began to express α-SMA, indicating the effective fibroblast activation and differentiation of part of these cells to myofibroblast.

**Figure 8.**
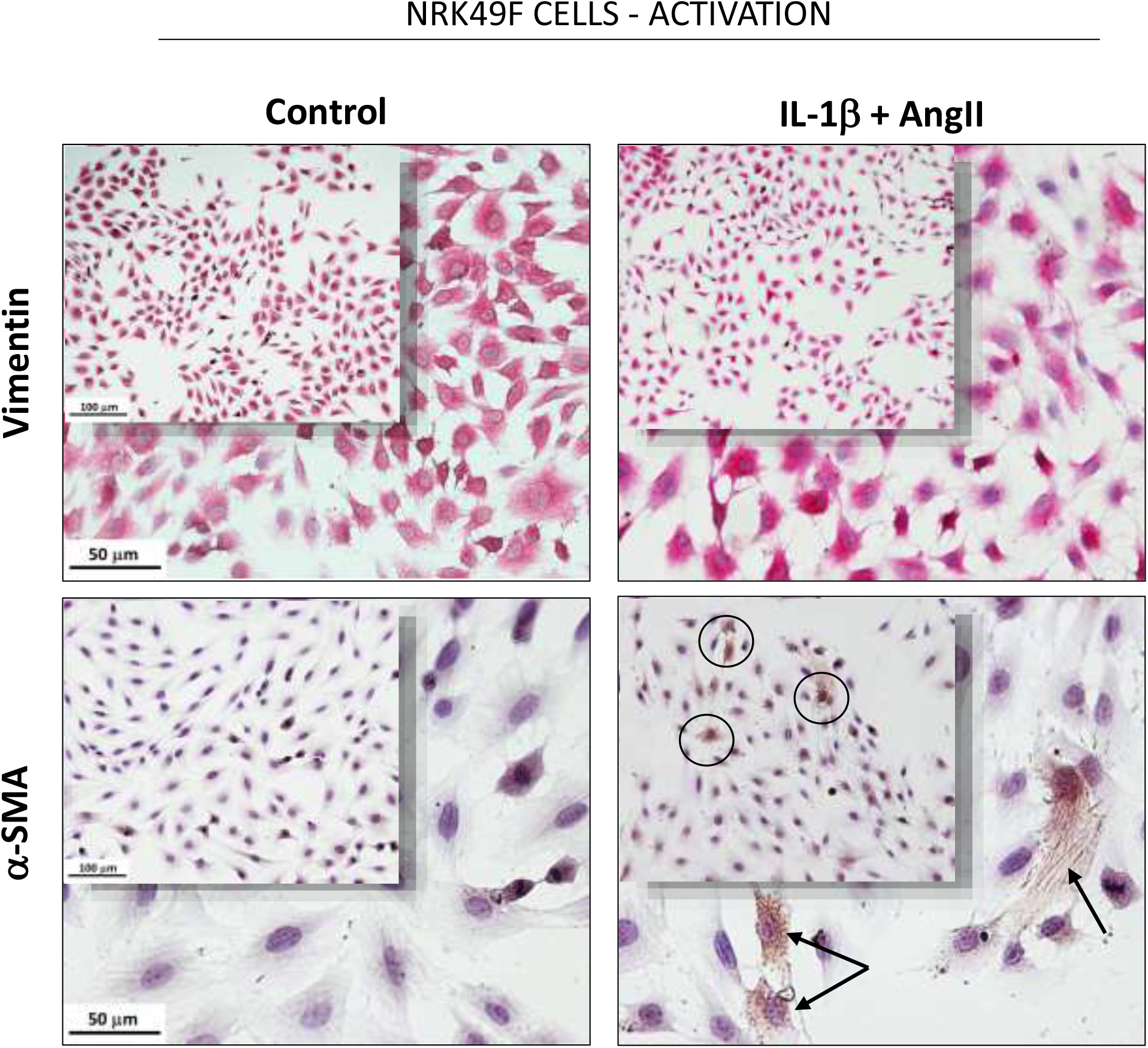
Representative micrographs of immunocytochemistry of untreated (Control) and IL-1β+AngII-stimulated NRK49F cells, stained for both a constitutive (vimentin) and a fibroblast activation-related (α-SMA) protein.

The results of RT-qPCR analysis of gene expression of pro and antifibrotic factors in NRK-49F cells are shown in **Figure 9**. IL-1β + AngII stimulus upregulated the expression of SAMD3 and SMAD7, as well as the expression of fibronectin, collagen I and collagen III in NRK-49F cells. While LOS treatment only reverted partially the overexpression of SMAD3, both TAM and associated LOS+TAM significantly normalized the expression of fibronectin, collagen I and collagen III, and reduced SMAD3 expression to levels lower than the observed in Control NRK49F cells. Additional illustrative immunocytochemistry for fibronectin and collagen I performed in cultured NRK-49F cells are shown in **Figure 10**, in which is it possible to verify that, untreated IL-1β + AngII-stimulated NRK-49F and LOS-treated cells exhibited exuberant positivity for fibronectin, suggesting fibronectin assembly and ECM overproduction, compared with the unstimulated NRK-49F or to the TAM and LOS+TAM-treated cells.

**Figure 9.**
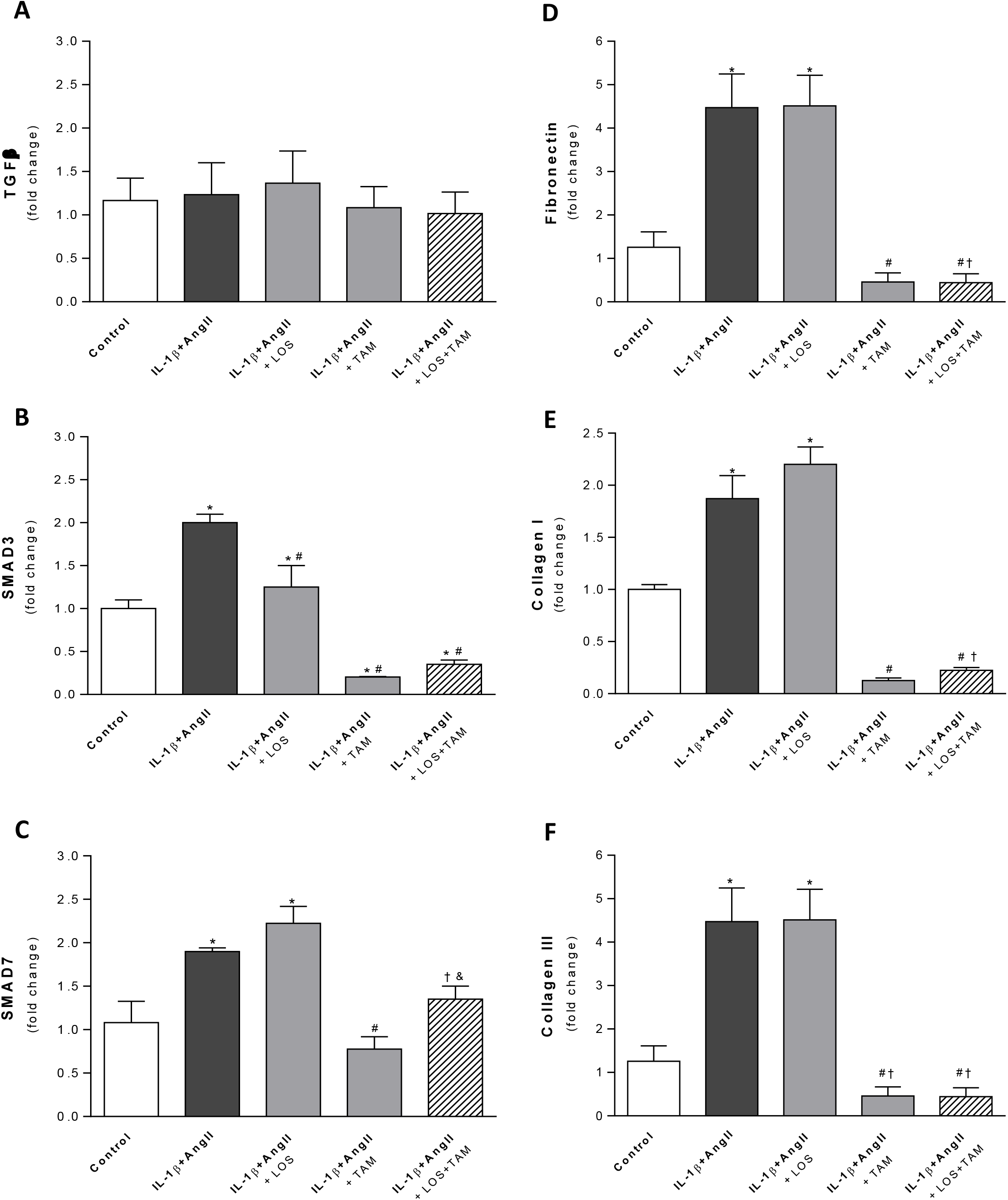
Quantitative RT-PCR of cultured NRK49F cells, submitted to the different treatments. RNA expression of pro and anti-fibrotic signaling factors; TGFβ **(A)**, SMAD3 **(B)** and SMAD7 **(C)**, as well as for the ECM proteins; collagen I **(D)** collagen III **(E)** and fibronectin **(F)**, were presented as bar graphs.

**Figure 10.**
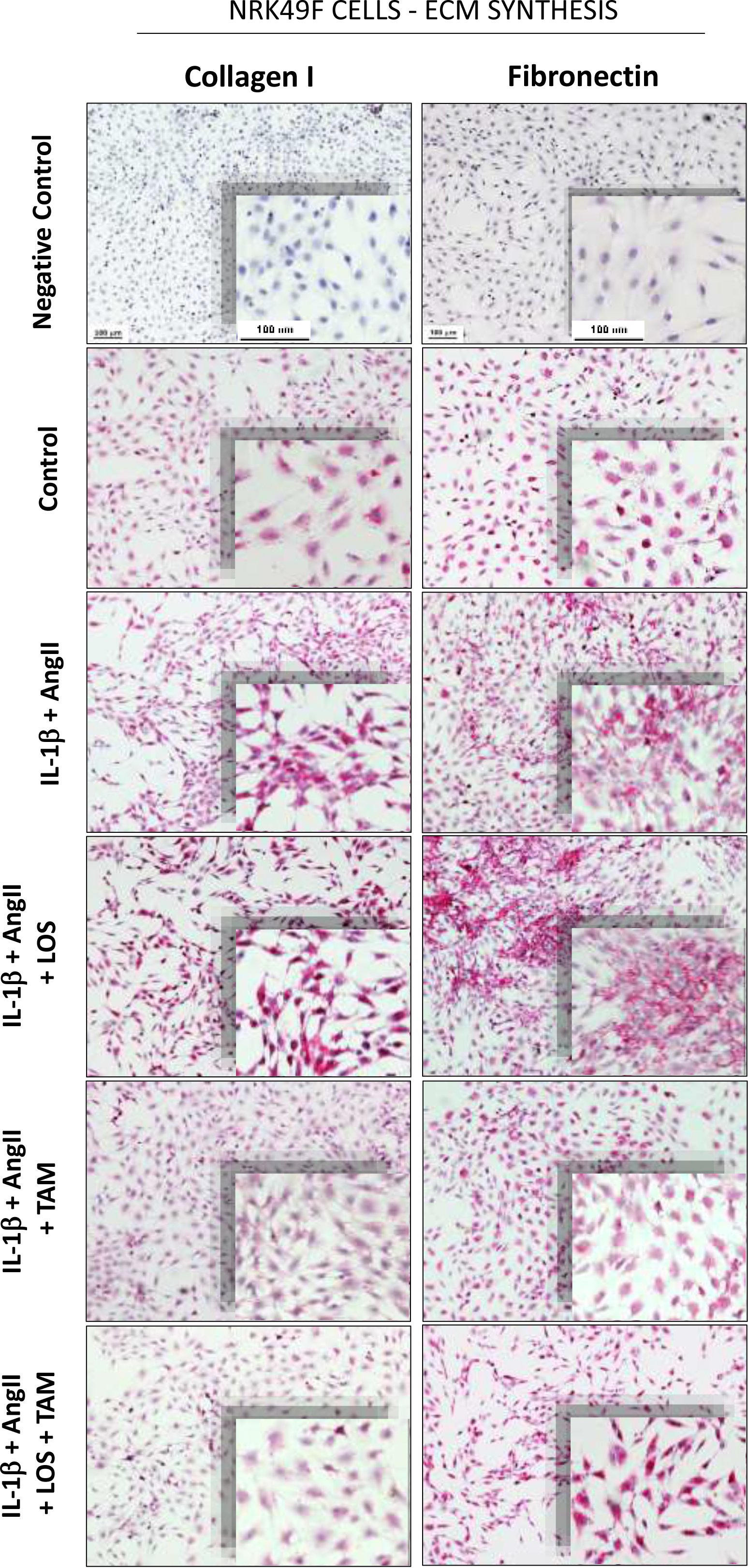
Representative micrographs of immunocytochemistry of NRK49F cells, stained for collagen I and fibronectin.

## DISCUSSION

In the present study we investigated the potential renoprotective effects of the therapeutic association of LOS+MMF+TAM on an experimental model of hypertensive nephrosclerosis, based on the chronic inhibition of NO synthesis, induced by L-NAME administration. The main aim of our research was to verify whether the combination of TAM, a selective estrogen receptor modulator, recommended for the treatment of positive estrogen receptor (ER+) breast cancer, to the currently employed conservative CKD treatment, here represented by RAAS blockade and immunosuppression, would promote additional anti-inflammatory and/or antifibrotic beneficial effects, when compared to the respective monotherapies.

Corroborating previous data, rats submitted to the L-NAME model of CKD developed severe hypertension, probably caused by glomerular and systemic vasoconstriction due to the lack of physiological vasodilatory effects of NO. Systemic hypertension and CKD are closely related conditions with an intricate cause/effect relationship. The decline of kidney function usually leads to high blood pressure, due to both a decreased ability of the kidneys to remove salt from the bloodstream, and an increased release of renal vasoconstrictive hormones. On the other hand, systemic hypertension sustained for long periods leads to the damage of multiple target organs, including the kidneys [7,8]. Hypertensive nephrosclerosis is one of the main causes of end-stage renal failure. High blood pressure also contributes to the aggravation of CKD, regardless of its etiology [1]. Accordingly, the clinical management of hypertension is currently one of the most employed strategies to control CKD progression [7,8]. According to our results, systemic hypertension induced by L-NAME administration was only partially reduced by the monotherapies with both LOS or MMF, and equally by the association of LOS+MMF+TAM. There was no synergistic effect of the combination of drugs as regards lowering blood pressure. The poor hemodynamic effect of therapeutic schemes may have limited the potential effects of the therapeutic association on the maintenance of renal function.

Along with the hypertension, NAME animals also exhibited a markedly increased urine albumin excretion rate, a clear evidence of renal impairment. Because of its strong predictive power for cardiovascular and renal events, albuminuria is one of the most important biomarkers of CKD progression, particularly in patients with hypertension or diabetes mellitus. Moreover, the reduction of albuminuria is the most important goal to prevent the progression of kidney disease in CKD patients. Usually, in this regard, significant benefits are achieved by the therapeutic treatment with RAAS inhibitors. According to our results, although all the tested monotherapies significantly limited the development of albuminuria in NAME animals, the antiproteinuric effect obtained with the combined treatment was noteworthy. Is spite of the severe sustained hypertension, LOS+MMF+TAM association completely averted the development of albuminuria in this CKD model.

The increased urinary albumin excretion is directly related to the disruption of one or more components of the glomerular filtration barrier, and with glomerular structural damage, generally caused or worsened by renal inflammation. Histological glomerular alterations are a common feature in most human and experimental nephropathies. Accordingly, severe glomerulosclerosis and glomerular collapse were observed in untreated NAME animals, and the association of LOS+MMF+TAM significantly prevented the development of glomerular histological damage, probably reflecting the effects of LOS (for glomerulosclerosis) and MMF (for collapsed glomeruli), thus evidencing that TAM did not exerted antagonism, blockade or inhibition upon the pharmacological effects of both LOS and MMF, and did not diminish the renoprotective effects observed with these drugs.

Kidney infiltration by inflammatory leukocytes has been demonstrated in a variety of non-immune mediated nephropathies, such as the hypertensive nephrosclerosis [4,19]. The recruitment of circulating monocytes, as well as the activation of resident renal macrophages often correlates positively with the worsening of renal function loss, in both human and experimental CKD [2]. Accordingly, in the present study, NAME animals showed exuberant renal inflammation, characterized by inordinate tubulointerstitial cell proliferation and massive infiltration of kidneys by both macrophages and lymphocytes. Surprisingly, similarly to the observed with LOS and MMF monotherapies, expected to exert inhibitory effects on macrophage and lymphocyte renal infiltration, as well as on interstitial proliferation rate, TAM monotherapy also exhibited independent significant anti-inflammatory proprieties. Therefore, the association of LOS+MMF+TAM seems to combine different mechanisms of action to abrogate renal inflammation. Inhibition of macrophage activity by TAM treatment has been demonstrated in both *in vitro* and *in vivo* studies, in which Tamoxifen promoted significant reduction in the transcription of important cell surface receptors, such as the fatty acid-binding proteins (FABPs) and the scavenger receptor class B member 3 (SCARB3 / CD36), which are involved in the monocyte/macrophage activation processes, and play a pivotal role in foam cell formation and in the development of atherosclerosis [25]. The anti-proliferative effects of RAAS blockade, associated with the reduction of IL1, IL6 and IL10 macrophage release, possibly achieved with the MMF treatment, may have boosted the anti-inflammatory effects of TAM. A synergetic effect among the tested drugs could also be plausible in this case. However, additional studies focused on the intracellular mechanisms of action of each employed drug and in the possible chemical interactions among them should be carried out to speculate this hypothesis.

Along with renal inflammation, the overproduction of ECM and the renal interstitial collagen accumulation are important histological features, commonly related to the worsening of CKD. In the present study renal cortical interstitial fibrosis, evidenced by the high percentage of Masson^+^ interstitial staining, myofibroblasts infiltration, as well as collagen I and fibronectin interstitial deposition, accompanied the progression of hypertensive nephrosclerosis in untreated NAME rats. All the tested therapies were effective in preventing renal fibrosis and α-SMA accumulation, while only TAM and the association of LOS+MMF+TAM significantly reduced collagen I and fibronectin accumulation in this CKD model, suggesting that TAM promoted additional antifibrotic effect to the therapeutic scheme, with no impairment of renoprotective action of LOS and MMF, when associated to these drugs. The suppressive effects of tamoxifen on fibrogenesis were first described in the early nineties, when Clark and collaborators described the drug to be effective and safe in the treatment of two patients with severe retroperitoneal fibrosis. Its effectiveness for the treatment of encapsulating peritoneal sclerosis, where than demonstrated, eight years later, by Allaria and co-authors. Based on these observations, Dellê and collaborators, from our research group, showed for the first time that TAM could exert protective effects on experimental progressive chronic kidney disease, in 2003. [26, 18, 27]. More recently, TAM was described to exert important antifibrotic effects in the experimental model of unilateral ureteral obstruction (UUO) in mice. Similarly, to the observed in the present study, TAM treatment reduced the production and deposition of ECM proteins in UUO kidneys, as well as the renal deposition of fibronectin and collagen [28, 29]. Although the exact mechanisms involved in the anti-inflammatory and antifibrotic effects of TAM are still poorly known, it exerts undoubted suppressive effects on fibroblast proliferation, activation and ECM secretion, evidenced in our *in vitro* results, which corroborate the current literature [30]. Since MMF, employed as an immunosuppressive drug in our *in vivo* protocol, is in fact a prodrug, which must be ingested and then metabolized into the pharmacologically active drug (mycophenolic acid), we were not able to perform *in vitro* studies with the full LOS+MMF+TAM association. However, we combined LOS to TAM is our analysis and clearly demonstrated that LOS did not impaired the suppressive effects of TAM on cultured fibroblasts, thus corroborating the idea that this drug combination may be safe and effective.

In summary, although further studies employing different CKD models are still required to confirm the efficacy and safety of the association of LOS+MMF+TAM, in the present paper we have provided evidence that this therapeutic scheme can be potentially useful to slow the progression of chronic nephropathy, since it lowered systolic blood pressure, prevented albuminuria, glomerular structural damage, and renal inflammation and promoted additional antifibrotic effect to the traditional conservative treatment of CKD, in the NAME model of hypertensive nephrosclerosis.

## CONCLUSION

Our pre-clinical observations suggested that the association of TAM to the conservative treatment of CKD, employing LOS and MMF, was safe and promoted additional renoprotective, anti-inflammatory and antifibrotic effect in a model of hypertensive nephrosclerosis in rats.

## Compliance with Ethical Standards

### Ethical Approval - Research involving animals

All the experimental procedures described in the present study were approved by the local Research Ethics Committee (CAPPesq, process No. 796/00) and were developed in strict conformity with our institutional guidelines and the international standards for manipulation and care of laboratory animals.

## Acknowledgments

The present research was financially supported by São Paulo Research Foundation (Process FAPESP - 03/05405-7).

## REFERENCES

[1] Romagnani P, Remuzzi G, Glassock R, Levin A, Jager KJ, et al. Chronic kidney disease. Nat Rev Dis Primers. 23(3), (2017):17088.

[2] Noronha IL, Fujihara CK, Zatz R. The inflammatory component in progressive renal disease - Are interventions possible? Nephrol Dial Transplant. 17(3), (2002):363–8.

[3] Zeisberg M, Neilson EG. Mechanisms of tubulointerstitial fibrosis. J Am Soc Nephrol. 21(11), (2010):1819–34.

[4] Fanelli C, Dellê H, Cavaglieri RC, Dominguez WV, Noronha IL. Gender Differences in the Progression of Experimental Chronic Kidney Disease Induced by Chronic Nitric Oxide Inhibition. Biomed Res Int. (2017):159739.

[5] Graciano ML, Cavaglieri RC, Dellê H, Casarini DE, Malheiros DMAC. Intrarenal reninangiotensin system is upregulated in experimental model of progressive renal disease induced by chronic inhibition of nitric oxide synthesis. J Am Soc Nephrol. 15(7), (2004):1805–15.

[6] Rüster C, Wolf G. Renin-angiotensin-aldosterone system and progression of renal disease. J Am Soc Nephrol. 17(11), (2006):2985–91

[7] Judd E, Calhoun DA. Management of Hypertension in CKD: Beyond the Guidelines. Adv Chronic Kidney Dis. 22(2), (2015):116–22.

[8] Townsend RR, Taler SJ. Management of hypertension in chronic kidney disease. Nat Rev Nephrol. 11(9), (2015):555–63.

[9] Arias SCA, Valente CP, Machado FG, Fanelli C, Origassa CS, et al. Regression of albuminuria and hypertension and arrest of severe renal injury by a losartan-hydrochlorothiazide association in a model of very advanced nephropathy. PLoS One. 2013:8(2), (2013):e56215.

[10] Fanelli C, Fernandes BHV, Machado FG, Okabe C, Malheiros DMAC, et al. Effects of losartan, in monotherapy or in association with hydrochlorothiazide, in chronic nephropathy resulting from losartan treatment during lactation. Am J Physiol Renal Physiol. 301(3), (2011):F580–7.

[11] Fujihara CK, Malheiros DMAC, Zatz R, Noronha IL. Mycophenolate mofetil attenuates renal injury in the rat remnant kidney. Kidney Int. 54(5), (1998):1510–9.

[12] Fujihara CK, Malheiros DMAC, Noronha IL, de Nucci G, Zatz R. Mycophenolate Mofetil Reduces Renal Injury in the Chronic Nitric Oxide Synthase Inhibition Model. Hypertension. 37(1), (2001):170–175.

[13] Utimura R, Fujihara CK, Mattar AL, Malheiros DMAC, Noronha IL, et al. Mycophenolate mofetil prevents the development of glomerular injury in experimental diabetes. Kidney Int. 63(1), (2003):209–16.

[14] Fujihara CK, Noronha IL, Malheiros DMAC, Antunes GR, de Oliveira IB, et al. Combined mycophenolate mofetil and losartan therapy arrests established injury in the remnant kidney. J Am Soc Nephrol. 11(2), (2000):283–90.

[15] van Bommel EFH, Pelkmans LG, van Damme H, Hendriksz TR. Long-term safety and efficacy of a tamoxifen-based treatment strategy for idiopathic retroperitoneal fibrosis. Eur J Intern Med. 24(5), (2013):444–50.

[16] Moustafellos P, Hadjianastassiou V, Roy D, Velzeboer NE, Maniakyn N, et al. Tamoxifen Therapy in Encapsulating Sclerosing Peritonitis in Patients After Kidney Transplantation. Transplant Proc. 38(9), (2006):2913–4.

[17] R. J. Loffeld, T. F. van Weel. Tamoxifen for retroperitoneal fibrosis. Lancet. 6:341(8841), (1993):382.

[18] Allaria PM, Giangrande A, Gandini E, Pisoni IB. Continuous ambulatory peritoneal dialysis and sclerosing encapsulating peritonitis: tamoxifen as a new therapeutic agent? J Nephrol. 12(6), (1999):395–7.

[19] Dellê H, Cavaglieri RC, Vieira Jr JM, Malheiros DMAC, Noronha IL. Antifibrotic effect of tamoxifen in a model of progressive renal disease. J Am Soc Nephrol. 23(1), (2012):37–48.

[20] Mancini GA, Carbonaro JF. Immunochemical quantitation of antigen by single radial immunodiffusion. Immunochemistry. 2(3), (1965):235–54.

[21] Jepsen LF, Mortensen PB. Interstitial fibrosis of the renal cortex in minimal change lesion and its correlation with renal function: a quantitative study. Virchows Arch A Pathol Anat Histol. 23;383(3), (1979):265–70.

[22] Wallenstein S, Zucker CL, Fleiss JL. Some statistical methods useful in circulation research. Circ Res. 47(1), (1980):1–9.

[23] Teles F, da Silva TM, da Cruz Jr FP, Honorato VH, de Oliveira Costa H, et al. (2015) Brazilian red propolis attenuates hypertension and renal damage in 5/6 renal ablation model. PLoS One. 21;10(1), (2015):e0116535.

[24] Fanelli C, Arias SCA, Machado FG, Okuma JK, Malheiros DMAC, et al. Innate And Adaptive Immunity are Progressively Activated in Parallel with Renal Injury in the 5/6 Renal Ablation Model. Sci Rep. 9;7(1), (2016):3192.

[25] Yu M, Jiang M, Chen Y, Zhang S, Zhang W, et al. Inhibition of Macrophage CD36 Expression and Cellular Oxidized Low Density Lipoprotein (oxLDL) Accumulation by Tamoxifen. J Biol Chem. 12;291(33), (2016):16977–89.

[26] C P Clark 1, D Vanderpool, J T Preskitt. The response of retroperitoneal fibrosis to tamoxifen Surgery. 109(4), (1991):502–6.

[27] Dellê H, Rocha JRC, Malheiros DMAC, Vieira JR JM, Noronha IL. Tamoxifen has protective effect preventing renal damage in chronic progressive renal disease. In: World Congress of Nephrology, 2003, Berlin. Nephrol Dial Transp 18, (2003):592.

[28] Kim CS, Kim IJ, Choi JS, Bae EH, Ma SK, et al. Tamoxifen ameliorates obstructive nephropathy through Src and the PI3K/Akt/mTOR pathway. Biol Cell. Jan;111(1), (2019):18–27.

[29] Tingskov SJ, Jensen MS, Pedersen CET, Araujo IBBA, Mutsaers HAM, et al. Tamoxifen attenuates renal fibrosis in human kidney slices and rats subjected to unilateral ureteral obstruction. Biomedicine & Pharmacotherapy, 133, (2021):111003.

[30] Silva FMO, Costalonga EC, Silva C, Carreira ACO, Gomes SA, et al. Tamoxifen and bone morphogenic protein-7 modulate fibrosis and inflammation in the peritoneal fibrosis model developed in uremic rats. Molecular Medicine 25, (2019):41.

